# Identification of mouse CD4^+^ T cell epitopes in SARS-CoV-2 BA.1 spike and nucleocapsid for use in peptide:MHCII tetramers

**DOI:** 10.1101/2023.11.16.566918

**Authors:** Laura Bricio Moreno, Juliana Barreto de Albuquerque, Jake M. Neary, Thao Nguyen, Kathryn M. Hastie, Sara Landeras-Bueno, Chitra Hariharan, Anusha Nathan, Matthew A. Getz, Alton C. Gayton, Ashok Khatri, Gaurav D. Gaiha, Erica Ollmann Saphire, Andrew D. Luster, James J. Moon

## Abstract

Understanding adaptive immunity against SARS-CoV-2 is a major requisite for the development of effective vaccines and treatments for COVID-19. CD4^+^ T cells play an integral role in this process primarily by generating antiviral cytokines and providing help to antibody-producing B cells. To empower detailed studies of SARS-CoV-2-specific CD4^+^ T cell responses in mouse models, we comprehensively mapped I-A^b^-restricted epitopes for the spike and nucleocapsid proteins of the BA.1 variant of concern via IFNγ ELISpot assay. This was followed by the generation of corresponding peptide:MHCII tetramer reagents to directly stain epitope-specific T cells. Using this rigorous validation strategy, we identified 6 reliably immunogenic epitopes in spike and 3 in nucleocapsid, all of which are conserved in the ancestral Wuhan strain. We also validated a previously identified epitope from Wuhan that is absent in BA.1. These epitopes and tetramers will be invaluable tools for SARS-CoV-2 antigen-specific CD4^+^ T cell studies in mice.

## Introduction

Severe acute respiratory syndrome coronavirus 2 (SARS-CoV-2) is the causative pathogen of the coronavirus disease 2019 (COVID-19) pandemic that has become an epic global health crisis (Li *et al*., 2021). The unprecedented effort to develop vaccines and treatments for this disease has put forth an urgency to better understand how adaptive immunity develops to the virus.

As with most other viral pathogens, effective immunity to SARS-CoV-2 involves a combination of humoral responses from B cells and antibodies as well as cellular responses from CD4^+^ and CD8^+^ T cells (Sette, Sidney and Crotty, 2023; Röltgen and Boyd, 2023). While the significance of virus-specific antibodies and particularly their neutralizing function in COVID-19 is unquestioned, there is growing appreciation for the parallel protective role provided by virus-specific T cells, due in part to their greater resistance to immune evasion by viral evolution and perhaps greater stability over time (Petrone *et al*., 2023).

T cells recognize peptides processed from antigenic proteins and presented by major histocompatibility complex (MHC) proteins on the surface of antigen-presenting cells (APC). Identification of the sequences of such peptide epitopes from SARS-CoV-2 proteins enables detailed analyses of SARS-CoV-2-specific T cell populations by informing the design of *in vitro* stimulation assays as well as the generation of peptide:MHC tetramer reagents that can directly stain SARS-CoV-2-specific T cells. Considerable progress has been made in the identification of human T cell epitopes for SARS-CoV-2 antigens, resulting in the generation and use of SARS-CoV-2-specific peptide:MHC tetramers for studies of viral epitope-specific T cell responses (Nelson *et al*., 2022). However, there are only a few reported sequences of mouse T cell epitopes that would benefit immunological studies in mouse models (Zhuang *et al*., 2021; Davenport *et al*., 2021; Joag *et al*., 2021; Wang *et al*., 2023; Yang *et al*., 2021; Smith *et al*., 2021).

In this report, we performed a comprehensive mapping of CD4^+^ T cell epitopes in SARS-CoV-2 spike and nucleocapsid proteins for C57BL/6 (B6) mice, the most widely used strain for immunological studies. Overlapping peptide libraries for these proteins were screened for reactivity to T cells in immunized mice by interferon-γ (IFNγ) ELISpot assay. Peptide:MHCII tetramer reagents were then generated for candidate peptide epitopes and used to measure the expansion of T cell populations with specificity for these epitopes and assess their immunogenicity. In all, we identified 6 novel epitopes in spike and 3 in nucleocapsid, which will be useful for future studies of T cell responses to SARS-CoV-2 in mice.

## Materials and methods

### Mice

C57BL/6 mice were purchased from Jackson Laboratory and housed under specific-pathogen-free conditions at Massachusetts General Hospital. Male and female mice were immunized between 6-16 weeks of age.

### Proteins

SARS-CoV-2 Wuhan-Hu-1 (D614G) spike was expressed in the “HexaPro” background (Hsieh *et al*., 2020) as previously reported (Hastie *et al*., 2021; Hastie *et al*., 2023). Briefly, ExpiCHO-S cells (Thermo Fisher) were maintained and transfected according to the manufacturer’s protocol. Transfected cells were harvested 7-8 days after transfection. Cultures were clarified by centrifugation and BioLock (IBA Life Sciences) added. The Twin-StrepII-Tagged-proteins were purified over a StrepTrap-HP column equilibrated with 25mM Tris pH 7.5, 200mM NaCl (TBS). After extensive washing, bound proteins were eluted in TBS buffer supplemented with 5mM d-desthiobiotin (Sigma Aldrich). Affinity-purified proteins were incubated with HRV-3C protease to remove purification tags and subsequently purified by size-exclusion-chromatography on a Superose 6 Increase column (GE Healthcare) in TBS

HexaPro SARS-CoV-2 BA.1 spike was expressed in ExpiCHO-S cells as described above or in *Drosophila* S2 cells as previously described for the production of peptide:MHCII tetramers (Moon and Pepper, 2018). Briefly, the same sequence was cloned into the pR expression vector and expressed in stably transfected S2 cells. Protein was harvested from cell culture supernatants and purified by binding to a nickel-nitrilotriacetic acid (Ni-NTA) column (Novagen) followed by elution with 1M imidazole containing 0.2% octyl β-D-glucopyranoside. After concentration through 100kD centrifugation filters (Millipore), the protein was further purified by size-exclusion chromatography on a Sephacryl 300 column (GE Healthcare) in PBS.

Recombinant SARS-CoV-2 BA.1 nucleocapsid protein was subcloned into the pET46 vector (Novagen) with an upstream 6xHis tag and expressed in Rosetta2 pLysS *E. coli* (Novagen). Cells were lysed with an M-110P microfluidizer (Microfluidics) in 50 mM HEPES pH 7.5, 500 mM NaCl, 10% glycerol, 20 mM imidazole, 6M urea, 1 ul Benzonase Nuclease (Millipore). Recombinant protein was purified by binding to Ni-NTA beads (Qiagen) and eluted in the same buffer supplemented with 300 mM imidazole. The resulting sample was then dialyzed overnight in Snakeskin dialysis tubing with 10-kDa pore size in refolding buffer (50 mM HEPES pH 7.5, 500 mM NaCl, 10% glycerol) and further purified by size-exclusion chromatography on a Superose 200 Increase column (GE Healthcare).

### Peptides

Peptide libraries for BA.1 spike and nucleocapsid (15-mers with 4aa overlap) were synthesized by the MGH Peptide Core (Naranbhai *et al*., 2022). Individual peptides to test minimal epitope sequences used in tetramers and native versions of mutated HexaPro spike sequences were custom ordered through Genscript.

### Immunizations

100 μg of recombinant protein or peptide was emulsified in a 1:1 mix of 0.9% saline and complete Freund’s adjuvant (CFA) and injected subcutaneously at the back of the neck and base of the tail (50 μl each site).

### T cell and APC isolation for ELISpot assays

The spleen and skin draining lymph nodes were harvested from mice 9-10 days postimmunization and processed into single cell suspensions by mechanical disruption over nylon mesh. CD4^+^ T cells were then isolated via a negative selection CD4^+^ T cell isolation kit (Miltenyi). For APCs, splenocytes were isolated from naïve mice, subjected to red cell lysis with ACK lysis buffer (Corning) at RT for 5 min, washed, and then irradiated at 2500 rads.

### ELISpot assays

IFNγ ELISpot assays were performed using a commercially available kit (Mabtech). 2.5 x 10^5^ CD4^+^ T cells were incubated with 2.5 x 10^5^ irradiated APCs in 200 μl R10 media (RPMI 1640 + 10% FBS, pen/strep, 2-ME) in each well of a 96-well pre-coated ELISpot plate. Single peptides from the spike and nucleocapsid libraries were added to each well at a final concentration of 20 μg/ml. Positive control wells were set up with 10 μg/ml Concanavalin A (ConA) and negative control wells with no stimulus. After overnight

(16-18h) incubation at 37C with 5% CO_2_, plates were washed and developed according to kit instructions. Dried plates were then analyzed on a Mabtech ASTOR ELISpot reader for quantification of spot-forming units (SFU), which were then expressed as SFU per 10^6^ CD4^+^ T cells after subtraction of background signal determined by negative controls.

### Tetramers

The generation of peptide:MHCII tetramers has been described in detail (Moon and Pepper, 2018). In brief, soluble heterodimeric I-A^b^ molecules covalently linked to peptide epitopes were expressed and biotinylated in stably transfected *Drosophila* S2 cells. Following immunoaffinity purification, these biotinylated peptide:MHCII complexes were titrated and tetramerized to streptavidin pre-conjugated to R-phycoerythrin (PE), allophycocyanin (APC), or R-phycoerythrin-cyanine-7 (PE-Cy7) fluorochromes (Prozyme, Thermo-Fisher).

### Tetramer-based T cell enrichment and flow cytometry

The spleen and skin draining lymph nodes were harvested from mice 7-8 days postimmunization and processed into single cell suspensions by mechanical disruption over nylon mesh. Tetramer-based enrichment of epitope-specific T cells from these samples was performed as described in detail (Legoux and Moon, 2012). Flow cytometry was performed with the Aurora spectral flow cytometer (Cytek). Data analysis was performed with FlowJo software (Treestar).

### Statistical analysis

All statistical analysis was performed on Prism (GraphPad).

## Results

### Screening of peptide libraries by IFNγ ELISpot

To facilitate detailed studies of SARS-CoV-2-specific CD4^+^ T cell responses in B6 mice, we performed a comprehensive screen of potential I-A^b^-restricted epitopes in SARS-CoV-2 spike and nucleocapsid proteins, which have been shown to be major immunogenic targets of CD4^+^ T cells during infection in humans (Grifoni *et al*., 2020). The sequences used in our study were from the Omicron (BA.1) variant strain due to the timing of when our studies began.

B6 mice were immunized subcutaneously with recombinant spike or nucleocapsid protein emulsified in complete Freund’s adjuvant (CFA), and 7-10 days later, expanded CD4^+^ T cells were isolated from pooled secondary lymphoid organs (spleen and skin-draining lymph nodes). To identify epitopes, the T cells were restimulated *in vitro* with individual peptides from a library of overlapping peptides (15-mer with 4 residue overlap) covering the entire sequence of each protein. Epitope-specific responses were gauged by the production of interferon γ (IFNγ) via ELISpot assay (Figure 1 and Supplemental File 1). In total,13 regions representing putative epitopes were identified in spike, and 3 were identified in nucleocapsid (Table 1).

**Table 1:**
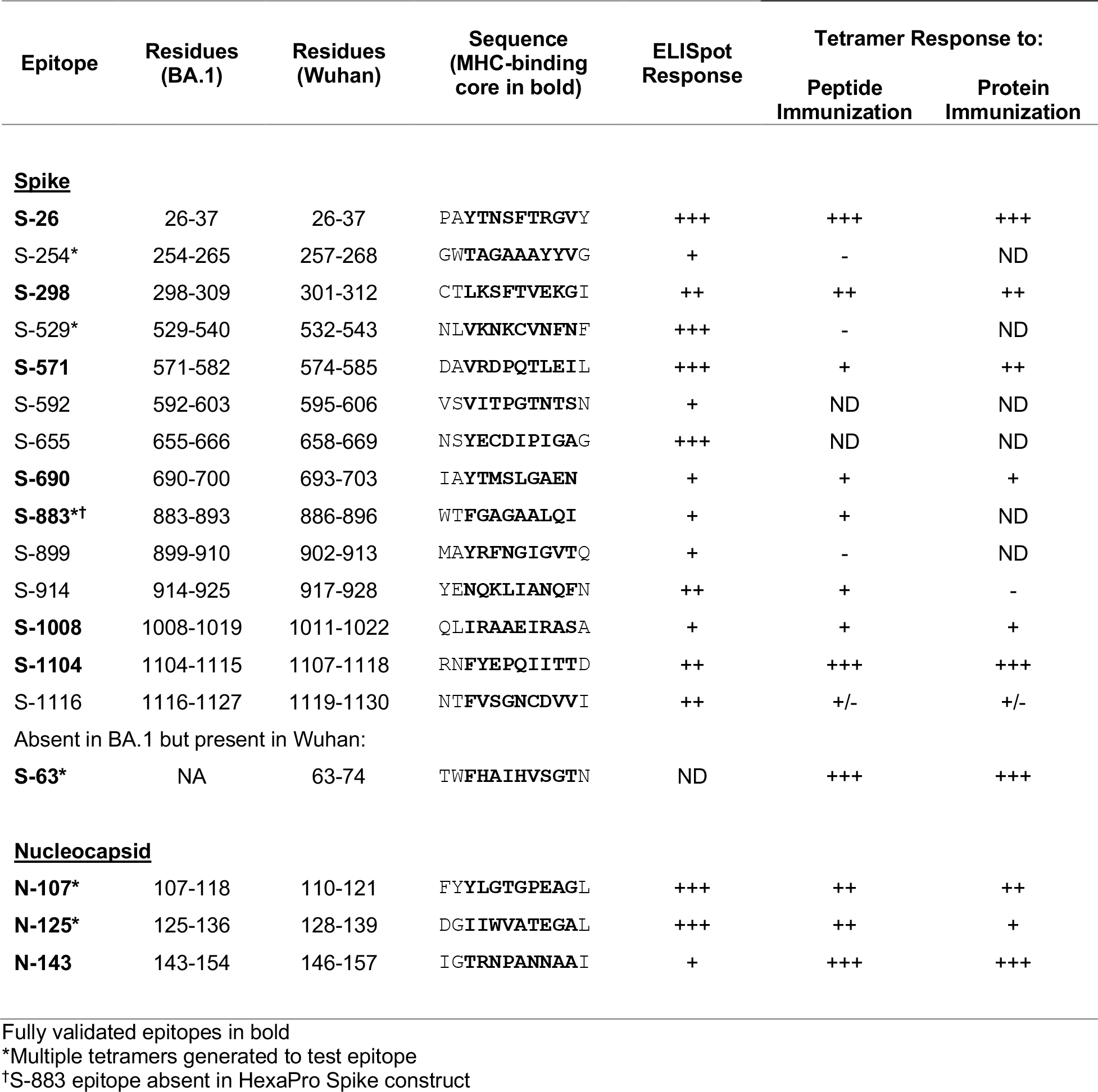
I-A^b^-restricted CD4^+^ T cell Epitopes in SARS-CoV-2 BA.1 Spike and Nucleocapsid

**Figure 1.**
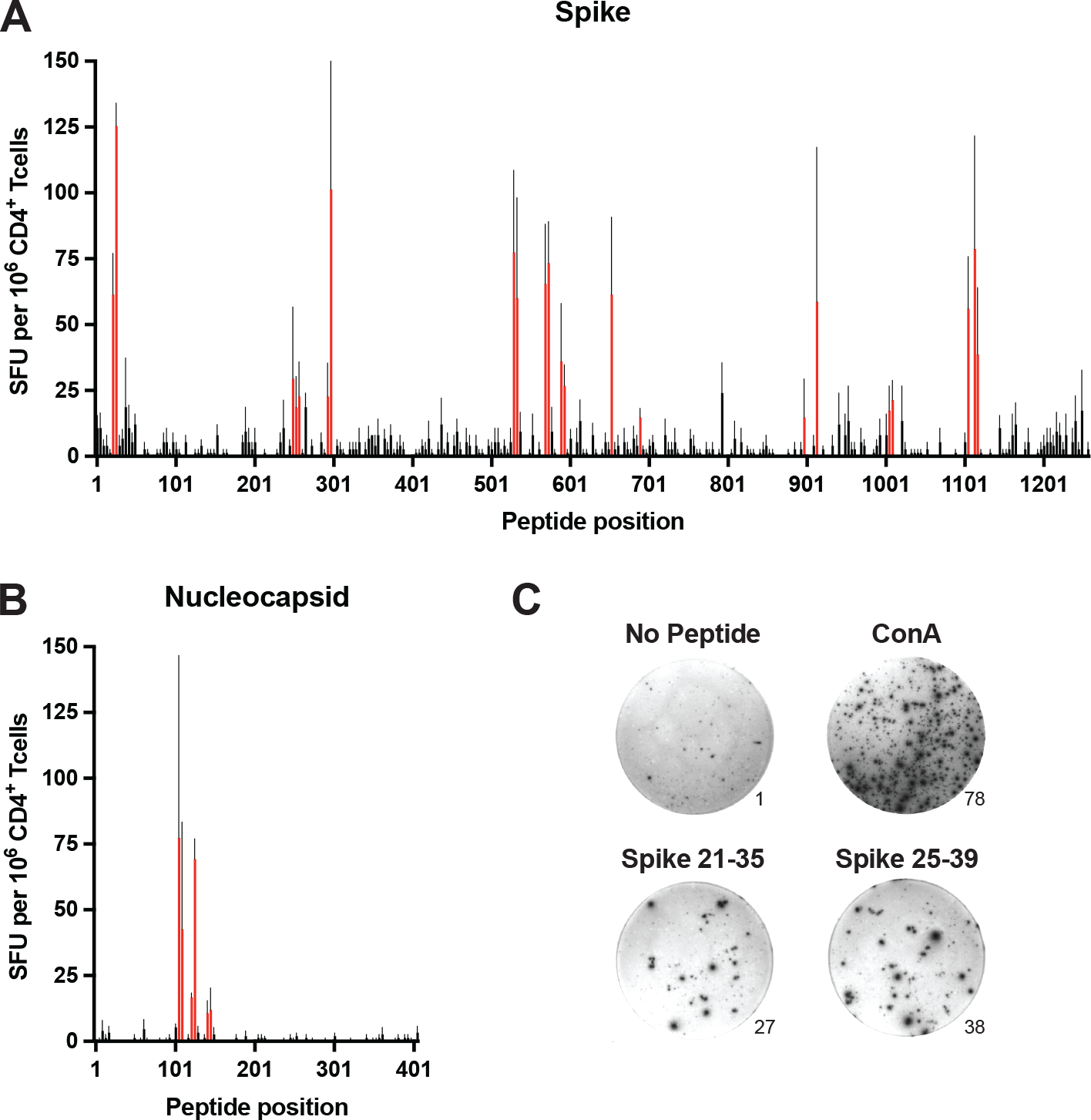
Comprehensive mapping of CD4^+^ T cell epitopes in SARS-CoV-2 BA.1 spike and nucleocapsid proteins. C57BL/6 mice were immunized subcutaneously (s.c.) with BA.1 spike or nucleocapsid proteins plus CFA as adjuvant, and 9-10 days later, CD4^+^ T cells from secondary lymphoid organs (SLO) were assessed for IFNγ production by ELISpot following *in vitro* restimulation with peptide libraries covering the entire sequence of **A)** spike and **B)** nucleocapsid. Mean values ± SEM are shown for n=3 independent experiments. Red bars indicate putative epitopes that were subsequently evaluated by peptide:MHCII tetramer staining. **C)** Representative images of ELISpot samples stimulated with PBS alone (No peptide, negative control), Concanavalin A (Con A, positive control), or overlapping peptides encompassing the S-26 epitope of spike (Spike 21-25 and Spike 25-39). Numbers at the lower right edges of images indicate the raw number of spotforming units counted per well.

### Tetramer-based validation of epitopes

To further validate these epitopes, we generated I-A^b^ tetramer reagents to directly stain epitope-specific CD4^+^ T cells. Tetramers were constructed with minimal peptide sequences (10-11 residues) representing each putative epitope as inferred from the overlap between neighboring immunogenic peptides as well as the use of computational tools that predict peptide binding registers in I-A^b^ (Lee *et al*., 2012; Dhanda *et al*., 2019) (Table 1). In some cases where minimal epitope sequences were not obvious, multiple tetramers were generated to cover the best possibilities from each immunogenic region (data not shown).

Mice were immunized with each minimal peptide, and 7 days later the expansion of CD4^+^ T cells specific for that peptide was assessed by corresponding tetramer staining and magnetic cell enrichment (Figure 2). Eight of the 13 spike epitopes and all three of the nucleocapsid epitopes exhibited potential immunogenicity based on observed increases in tetramer-positive T cell frequencies following immunization.

**Figure 2.**
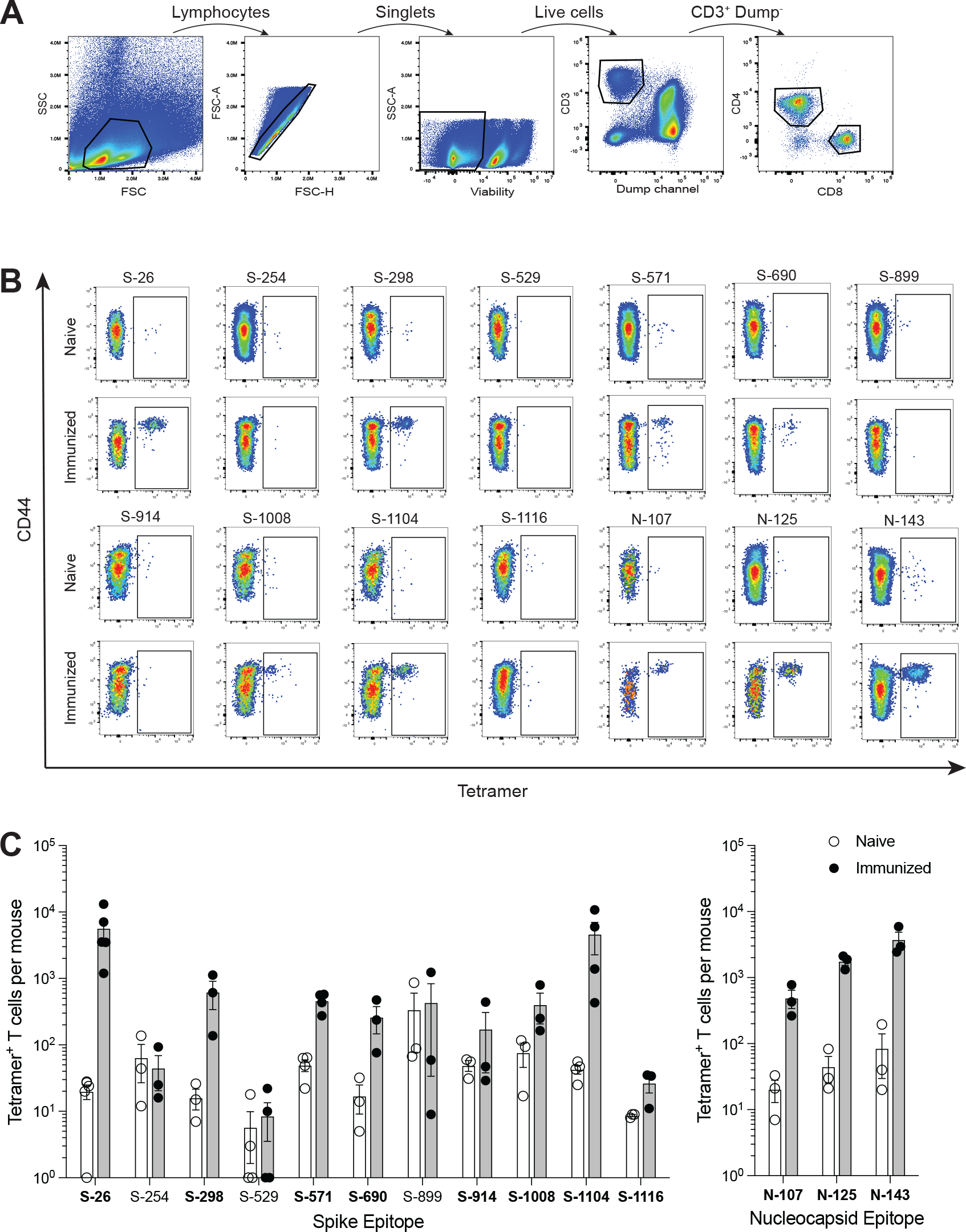
Identification of SARS-CoV-2 epitope-specific CD4^+^ T cells by peptide:MHCII tetramers. C57BL/6 mice were immunized s.c. with candidate epitope peptides plus CFA as adjuvant, and 7 days later, epitope-specific CD4^+^ T cells from the SLO were detected by peptide:MHCII tetramer-based cell enrichment and flow cytometry. **A)** Flow cytometry gating strategy for analysis of lymphocyte^+^ single cell^+^ live^+^ dump (B220, CD11b, CD11c, F4/80)^−^ CD3^+^ CD4^+^ events. **B)** Representative flow cytometry plots of CD4^+^ gated events illustrating tetramer staining of epitope-specific T cells from naïve and corresponding peptide-immunized mice. **C)** Quantification of epitope-specific CD4^+^ T cells from naïve and immunized mice. Mean values ± SEM are shown for n=3-5 mice per epitope across multiple independent experiments. Bold font indicates epitopes subsequently evaluated in protein immunization experiments.

Despite having excellent MHC-binding prediction scores, tetramers for spike epitopes S-592 and S-655 were consistently fraught with issues of high background staining and the immunogenicity of these epitopes could not be ascertained (data not shown).

Peptides were chosen for these immunizations to provide maximal levels of peptide:MHCII complex presentation that could test whether the epitope was capable of evoking a T cell response. Once epitopes were validated in this way, tetramers were then used to detect epitope-specific T cells following immunization of mice with whole spike or nucleocapsid protein (Figure 3). This additional step addressed the immunogenicity of each epitope when the efficiency by which it is processed from protein and presented by antigen-presenting cells is also considered. All but two of the 8 epitopes validated by peptide immunization elicited some level of response by protein immunization. Responses ranged from minimal to robust, and they reflected the same general hierarchy of epitope immunodominance seen in the peptide studies.

**Figure 3.**
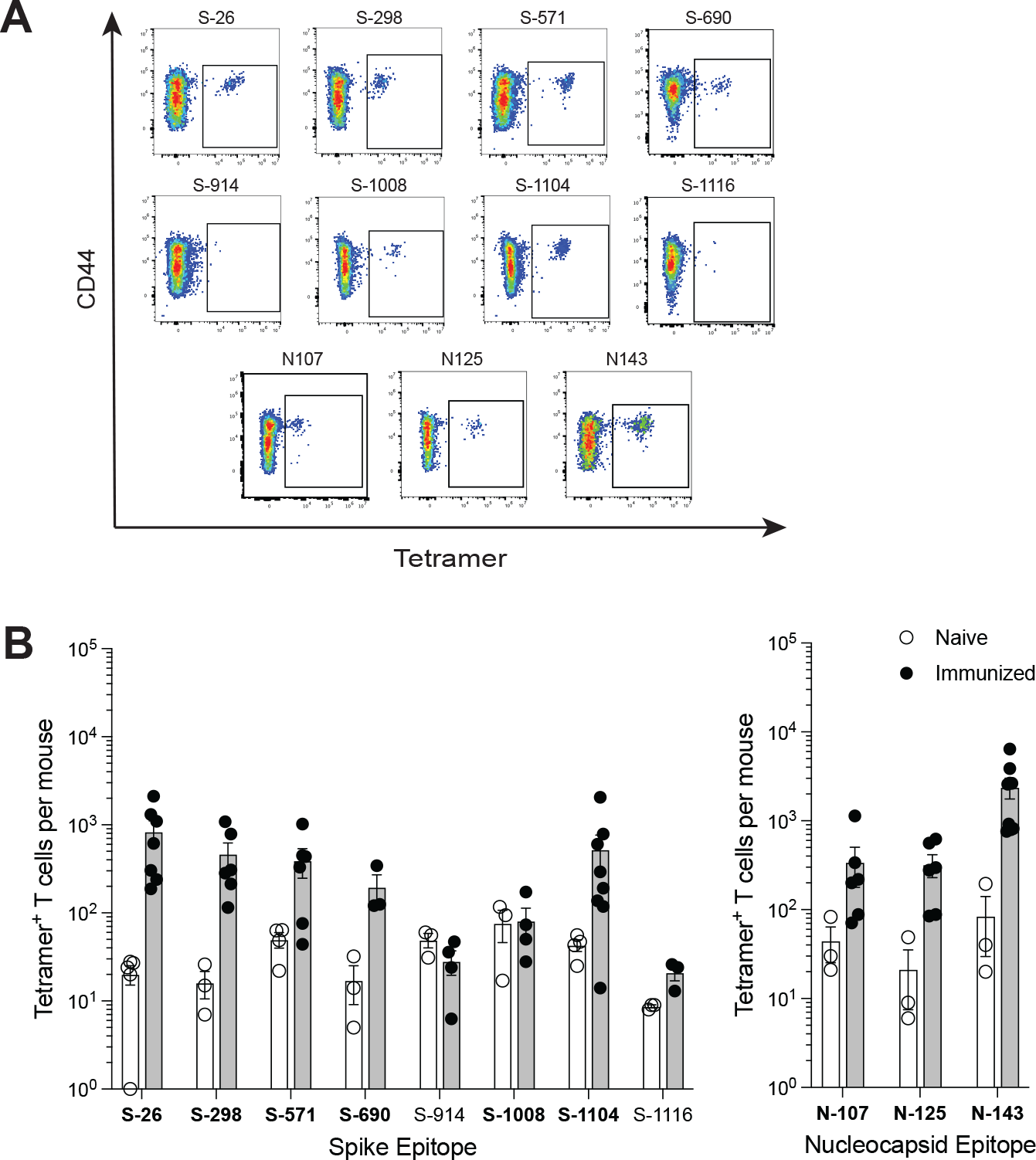
Comparison of SARS-CoV-2 epitope-specific CD4^+^ T cell responses after protein immunization. C57BL/6 mice were immunized s.c. with spike or nucleocapsid protein plus CFA as adjuvant, and 7 days later, epitope-specific CD4^+^ T cells from the SLO were detected by tetramer-based cell enrichment and flow cytometry. **A)** Representative flow cytometry plots of CD4^+^ gated events illustrating tetramer staining of epitope-specific T cells from protein-immunized mice. **B)** Quantification of epitope-specific CD4^+^ T cells from naïve and immunized mice. Mean values ± SEM are shown for n=3-9 mice per epitope across multiple independent experiments. Bold font indicates epitopes validated with increases in frequency upon immunization.

### Validation of additional epitopes

The spike protein used in our studies was the previously described HexaPro construct engineered to remain in a prefusion trimer complex due to the introduction of 6 stabilizing proline substitutions (Hsieh *et al*., 2020). To determine whether any potential epitopes were affected by these mutations, we immunized mice with peptides corresponding to native sequences at these locations and assessed T cell responses by IFNγ ELISpot (Figure S1). One of these peptides generated a weak response which was nonetheless validated by tetramer staining of expanded T cells in mice immunized with a minimal epitope sequence from this region (S-883, Table 1).

All of the identified SARS-CoV-2 BA.1 spike and nucleocapsid epitopes in our study are completely conserved from the ancestral Wuhan-Hu-1 strain. However, a previously reported immunodominant epitope (spike residues 62-78) in Wuhan (Zhuang *et al*., 2021) is mutated in BA.1 and therefore was not identified in our screen. To compare the immunogenicity of this epitope to the new ones identified in our current study, we generated a corresponding tetramer with a minimal epitope sequence (S-63, Table 1) and tested it in mice immunized with peptide or Wuhan spike protein (Figure S2). Responses to this epitope were very robust and superior to the responses for any of the newly identified epitopes.

## Discussion

In this study, we provide a comprehensive mapping and evaluation of mouse I-A^b^-restricted CD4^+^ T cell epitopes in spike and nucleocapsid, two of the major immunogens of the SARS-CoV-2 virus. This was done systematically by screening a library of overlapping peptides covering the entire sequences of these proteins by IFNγ ELISpot assay. This was followed up with the generation of peptide:MHCII tetramer reagents, which were used to validate candidate epitopes as well as evaluate their relative immunogenicity The result of our labor-intensive empirical approach was a more rigorous identification of epitopes than what computational approaches of epitope prediction alone can provide. However, we did use computational approaches to help narrow down minimal peptide epitope sequences during the design of tetramers.

In total, we discovered 6 reliably immunogenic epitopes in spike (S-26, S-298, S-571, 6-690, S-1008, and S-1104) and 3 in nucleocapsid (N-107, N-125, N-143) (summarized in Table 1), which to the best of our knowledge are all novel. One additional weak spike epitope (S-883) lies in a region affected by a proline substitution (A889P) used in the prefusion stabilization of the HexaPro construct (but not current vaccines in the U.S.), highlighting potential caveats of using Hexapro and other engineered versions of viral proteins for immunological studies.

We also validated a previously identified immunodominant spike epitope (S-63) that is present in the Wuhan strain but absent in the BA.1 variant that was studied in our report. A tetramer representing this epitope (residues 62-76) was successfully used in a previous study (Mao *et al*., 2022). The tetramer created in our current study differs by containing a smaller, minimal peptide sequence (residues 63-74) for this epitope. There is another weaker previously identified epitope in nucleocapsid (residues 9-23) of Wuhan that is also absent in BA.1, but we did not investigate this (Zhuang *et al*., 2021). Interestingly, the previous studies mapping CD4^+^ T cell epitopes in C57BL/6 mice only identified S-63 and not any of our novel epitopes (Zhuang *et al*., 2021; Wang *et al*., 2023). As the S-63 epitope is considerably more immunodominant than the others, this discrepancy could simply be due to differences in the sensitivity of our methods, or perhaps differences in epitope immunodominance patterns resulting from the different ways in which mice were immunized with antigen.

The most notable limitation of our study is that despite our comprehensive efforts, additional epitopes may have been missed. Several candidates identified by the ELISpot assay were not successfully validated by tetramers. It is not clear whether this was due to inaccurate prediction of minimal epitope sequences that were used in tetramer design or other technical issues related to tetramer generation. Although we screened potential epitopes masked by proline substitutions in HexaPro spike, we did not screen the truncated 63 amino acids of the C-terminus that are replaced with a Foldon domain in this construct (Hsieh *et al*., 2020). Finally, our assay relies specifically on T cell production of IFNγ, and although unlikely, it is possible that other epitopes preferentially generating other non-Th1 cell lineages were missed.

All of the newly identified epitopes are conserved between BA.1 and the ancestral Wuhan strain as well as Alpha, Gamma, Delta, BA.2, BA.5, XBB.1.5, EG.5.1, and the mouse adapted strain MA10 (Leist *et al*., 2020), and differences with Beta, BA.4, and BQ.1.1 are limited to just one epitope each (S-690, N-143, N-125, respectively). The S-63 epitope notwithstanding, this is not entirely surprising as there is unlikely to be immune selective pressure against mouse T cell epitopes in a human virus. However, it is noteworthy that human T cell epitopes have also been generally conserved across variants of concern, suggesting that immune evasion is driven predominantly by antibody rather than T cell recognition (Tarke *et al*., 2021; Naranbhai *et al*., 2022). A welcome consequence of this conservation is that the epitopes described here for BA.1 can also be used for studies of Wuhan and other variant strains. It will be interesting to see if further epitope mapping for Wuhan and other variants will identify additional epitopes discordant with BA.1.

The identification of these I-A^b^-restricted epitopes will be very useful for studies of CD4^+^ T cell immunity in C57BL/6 mice, the most commonly used strain in immunological research and the background for most hACE2 transgenic mice used in SARS-CoV-2 infection studies (McCray *et al*., 2007; Muñoz-Fontela *et al*., 2020). There is an appreciable difference in immunogenicity between the two strongest epitopes (S-26, S-1104) and the rest, so we propose the use of these epitopes along with S-63, if applicable, for robust studies of antigen-specific CD4^+^ T cells, especially if tetramers are to be used.

## Supporting information

Supplementary File 1 - ELISpot counts

## Ethics statement

All experiments were approved by the Institutional Animal Care and Use Committee (IACUC) of Massachusetts General Hospital, an American Association for the Accreditation of Laboratory Animal Care (AAALAC)-accredited animal management program.

## Author contributions

LBM, JBA, AL, and JM conceived the study. LBM, JBA, JN, and TN performed the majority of the experiments. JN, KM, SLB, CH, JM, and EOS generated recombinant proteins. AN, MG, AG, AK, and GG generated peptide libraries. LBM, JBA, AL, and JM designed the experiments and analyzed the data. JBA and JM wrote the manuscript. All authors reviewed and approved the final manuscript.

## Acknowledgments

We thank Lucy Kuhn and Tianyi Bai for assistance with tetramer and protein production, and Daniel Shin, Ryan Nelson, and Maisa Takenaka for helpful comments.

## Supplementary materials

The Supplementary Material for this article can be found online at:

