## Supplementary Material for "Identification of mouse CD4^+^ T cell epitopes in SARS-CoV-2 BA.1 spike and nucleocapsid for use in peptide:MHCII tetramers"

### Figure S1

**A**

| Peptide | Sequence |
|---|---|
| 806-822 | PSKPSKRSFIEDLLFNKV |
| 881-904 | SGWTFGAGAALQIPFAMQMAYRFN |
| 931-947 | IQDSLSSTASALGKLQD |
| 975-992 | NDIFSRLDKVEAEVQIDR |

**B**

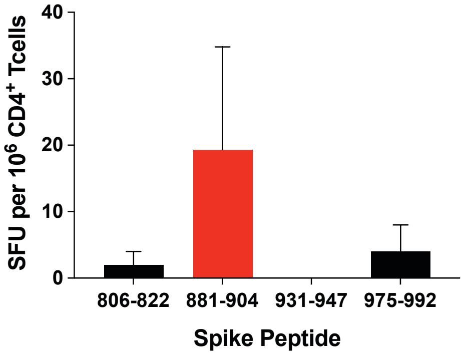


**C**

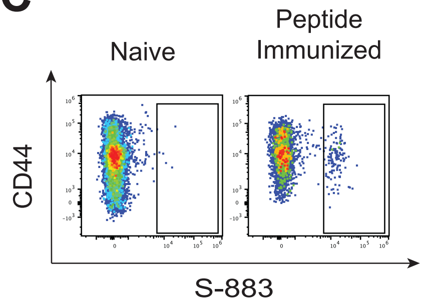


**D**

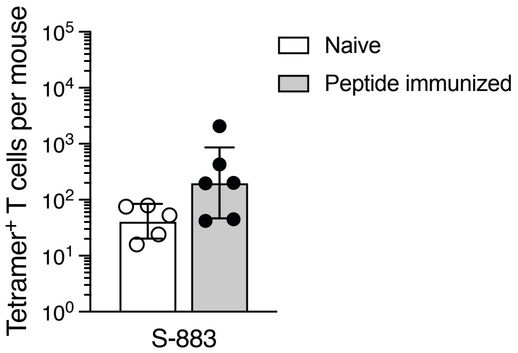


**Figure S1. Identification of lost epitopes in Hexapro spike**

**A)** Peptides were generated covering the regions of spike that were affected by proline substitutions in the Hexapro construct. Native residues that were replaced by prolines are indicated in red. **B)** C57BL/6 mice were immunized s.c. with a mix of the 4 peptides plus CFA as adjuvant and 9-10 days later, CD4$^+$ T cells were tested for reactivity to each individual peptide by IFN$\gamma$ ELISpot assay. Mean values ± SEM are shown for n=2-6 mice processed across multiple independent experiments. **C)** Representative flow cytometry plots of CD4$^+$ gated events illustrating S-883 tetramer staining of epitope-specific T cells from naïve and S-883 peptide-immunized mice. **D)** Quantification of S-883-specific CD4$^+$ T cells from naïve and peptide-immunized mice. Mean values ± SEM are shown for n=5-6 mice per epitope across multiple independent experiments.

### Figure S2

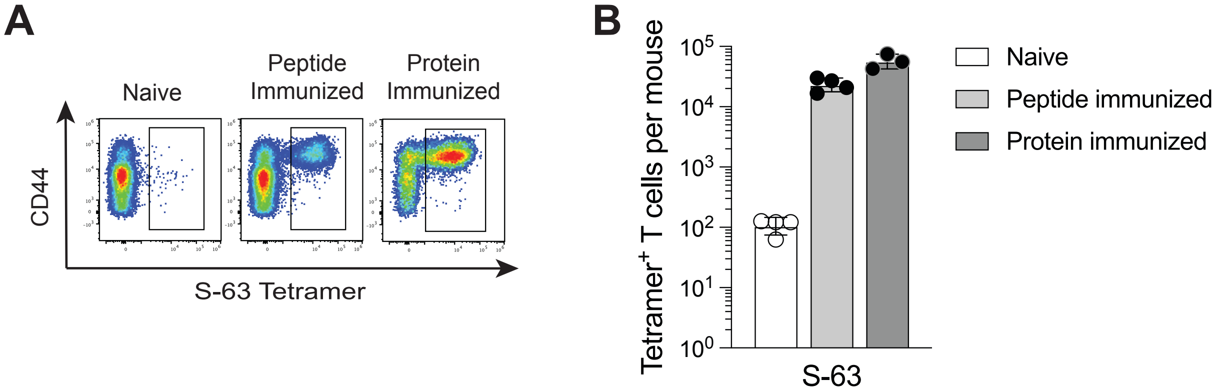


**Figure S2. Tetramer analysis of CD4⁺ T cells specific for the S-63 epitope present in the ancestral strain of SARS-CoV-2, but not BA.1**

**A)** Representative flow cytometry plots of $CD4^+$ gated events illustrating S-63 tetramer staining of epitope-specific T cells from naïve, S-63 peptide-immunized, or Wuhan D614G spike protein-immunized mice. **B)** Quantification of S-63-specific $CD4^+$ T cells from naïve, peptide-immunized, and protein-immunized mice. Mean values ± SEM are shown for n=3-4 mice per epitope across multiple independent experiments.
